# Considering decoupled phenotypic diversification between ontogenetic phases in macroevolution: An example using Triggerfishes (Balistidae)

**DOI:** 10.1101/2022.01.11.475856

**Authors:** Alex Dornburg, Katerina L. Zapfe, Rachel Williams, Michael E. Alfaro, Richard Morris, Haruka Adachi, Joseph Flores, Francesco Santini, Thomas J. Near, Bruno Frédérich

**Affiliations:** University of North Carolina at Charlotte, Charlotte, NC, USA; School of Natural Sciences, Bangor University, UK; Department of Ecology and Evolutionary Biology, University of California, Los Angeles, CA, USA; North Carolina Museum of Natural Sciences, Raleigh, NC, USA; Associazione Italiana per lo Studio della Biodiversità, Pisa, 56100, Italy; Department of Ecology and Evolutionary Biology, Yale University, New Haven, CT, USA; Laboratory of Functional and Evolutionary Morphology, FOCUS, University of Liège, Liège, Belgium

**Author notes:** Corresponding authors: Alex Dornburg, Bruno Frédérich. **Author Contributions** AD, RW, FS, BF, and KLZ conceived of the study. AD, KLZ, HA, JF, and RW collected data. AD, KLZ, BF performed analyses. AD and BF wrote the initial draft of the manuscript. All other authors contributed to the writing and subsequent development of the manuscript. **Data availability** Data is publicly available. Supplemental tables are included with this submission and alignments, phylogenies, and code are available on Zenodo: DOI: 10.5281/zenodo.5838141.

**Keywords:** Adaptive Radiation, Functional Morphology, Ecology, Nursery habitat, Ontogeny

## Abstract

Across the Tree of Life, most studies of phenotypic disparity and diversification have been restricted to adult organisms. However, many lineages have distinct ontogenetic phases that do not reflect the same traits as their adult forms. Non-adult disparity patterns are particularly important to consider for coastal ray-finned fishes, which often have juvenile phases with distinct phenotypes. These juvenile forms are often associated with sheltered nursery environments, with phenotypic shifts between adults and juvenile stages that are readily apparent in locomotor morphology. However, whether this ontogenetic variation in locomotor morphology reflects a decoupling of diversification dynamics between life stages remains unknown. Here we investigate the evolutionary dynamics of locomotor morphology between adult and juvenile triggerfishes. Integrating a time-calibrated phylogenetic framework with geometric morphometric approaches and measurement data of fin aspect ratio and incidence, we reveal a mismatch between morphospace occupancy, the evolution of morphological disparity, and the tempo of trait evolution between life stages. Collectively, our results illuminate how the heterogeneity of morpho-functional adaptations can decouple the mode and tempo of morphological diversification between ontogenetic stages.

## Introduction

For over a century, ecological opportunity has been regarded as a fundamental feature driving patterns of evolutionary diversification (Cope 1871; Simpson 1945; Bock 1965; Liem 1973; Hunter and Jernvall 1995; Schluter 2000; Price 2010; Dumont et al. 2012). However, research assessing the relationship between ecological opportunity and macroevolutionary dynamics is often limited to adult organisms (Alfaro et al. 2005; Losos 2009; Dornburg et al. 2011; Near et al. 2012; Frédérich et al. 2016; Wainwright and Price 2016; Cooney et al. 2017), with larvae and juveniles remaining largely neglected until recently (Sherrat et al. 2017; Valero et al. 2017). Such an ontogenetically restricted perspective is problematic as changes in one or more aspects of an organism’s niche during ontogeny (i.e. ecophases) are common across the Tree of Life, with examples documented in various lineages such as reptiles (Mushinsky et al. 1982); archosaurs (Marchetti and Price 1989; Subalusky et al. 2009); insects (Dopman et al. 2002); plants (Parish and Bazzaz 1985; Miriti 2006; Yang and Rudolf 2010); and fishes (Gagliano et al. 2007; Frédérich et al. 2008; Frédérich et al. 2010; Kimirei et al. 2013) to name but a few. Acknowledging that ontogenetic niche shifts might act as a potential source of ecological opportunity, a narrowed focus on only a single life-stage of a lineage may obscure major drivers of ecomorphological diversity. In the worst case, not considering different life stages of an organism in macroevolutionary studies can lead to either biased or incomplete conclusions regarding the general rules governing organismal evolution. Developing an expanded ontogenetic perspective on patterns of phenotypic diversification therefore represents an oft-neglected, yet critical aspect of evolutionary biology.

The observation that selective pressures are heterogeneous between ecophases has been discussed since the last century (Istock 1967; Wilbur 1980; Gagliano et al. 2007). Recent investigations of the transition between aquatic and terrestrial life stages in the complex life cycle of amphibians have illustrated a decoupling in the mode of morphological evolution across ecophases (Sherrat et al. 2017; Valero et al. 2017), highlighting the role of complex life cycles in shaping also the rate of morphological evolution (Fabre et al. 2020). Although it is clear that ontogenetic niche shifts can promote variation in the pattern of phenotypic diversification between life stages, this perspective has rarely been considered in the macroevolutionary history of marine fishes–an assemblage that represents a quarter of all living vertebrates. Many marine teleost fishes that occupy open or high flow environments only do so during their adult stage (Fulton and Bellwood 2002). In contrast, their juvenile forms generally inhabit nursery habitats that are characterized by three-dimensional complexity and shelter from changes in flow regime and predation (Nagelkerken et al. 2000). Across coastal environments, shifts in habitat usage throughout ontogeny are common (Lecchini and Galzin 2005; Amorim et al. 2018; Moussa et al. 2020) and subject transitioning individuals to new hydrodynamic regimes and ecological opportunities. For locomotor associated traits, these ontogenetic niche transitions are likely to place many aspects of juvenile and adult stage phenotypes under opposing selective pressures. For example, body or fin shapes differently affect acceleration, sustained swimming or mobility (Vogel 2008), often reflecting a functional trade-off between locomotor optimization for maneuverability versus prolonged cruising (Fossati 2009). Such a physical trade-off coupled with variation in flow regimes, resource availability, and predation risks is likely to have decoupled the tempo and the mode of locomotor phenotype diversification between ecophases of numerous marine fish lineages. Given that fin and body shapes are commonly used as representative phenotypes in macroevolutionary investigations of marine fishes, broadening our perspective beyond a single ecophase is therefore necessary to expand our knowledge of how marine fish diversify.

Triggerfishes (Balistidae) provide an ideal case study from which to investigate the relationship between juvenile and adult stages and the diversification of locomotor morphology. These iconic reef fishes possess a distinct mode of locomotion, balistiform locomotion, that couples oscillations and undulations of the dorsal and anal fins as the primary axis of forward propulsion (Lighthill et al. 1990; Sfakiotakis et al. 1999). The evolutionary novelty of balistiform locomotion is thought to have driven the adaptive evolution of changes in fin and body shapes between high and low flow regime associated species (Dornburg et al. 2011), as well as repeated convergence in habitat-associated body forms and fin shapes (Dornburg et al. 2008, 2011; Santini et al. 2013; McCord and Westneat 2016a). However, this hypothesis is based on sampling restricted to adult specimens. Triggerfishes possess a distinct juvenile form relative to their adult stage that is characterized by a more discoid body shape and the lack of high aspect ratio fins (Matsuura and Katsuragawa 1981). This condition is reminiscent of adult phenotypes in low-flow reef species such as the humuhumu (*Rhinecanthus rectangulus*) that possess deep bodies and long low aspect ratio dorsal and anal fins. In contrast, adult forms of high-flow open-ocean species such as the white-spotted oceanic trigger (*Canthidermis maculatus*) are characterized by shallow bodies with high aspect ratio dorsal and anal fins (Dornburg et al. 2011). The morphological variation associated with flow regimes is likely associated with habitat shifts that occur during the growth of different species. For example, in Moorea lagoon (French Polynesia), the juveniles of the white-banded triggerfish (*Rhinecanthus aculeatus*) live in sandy areas close to the beach while adults are found on the barrier reef in areas with high depth, strong current and high cover of living coral (Lecchini & Galzin 2005). The presence of ecophases as well as ontogenetic variation in functional morphology of swimming apparatus (i.e. body and fins) provides the opportunity to test if the evolutionary novelty of balistiform locomotion that is hypothesized to have catalyzed rapid morphological evolution early in the history of adult triggerfishes had a similar effect on the juvenile ecophase.

Here we investigated changes in locomotor morphology between adult and juvenile triggerfish phases across a time-calibrated phylogeny. We used geometric morphometrics to quantify median fin and body shapes, and also quantified fin aspect ratios from all major lineages. As changes in aspect ratio additionally require changes in the angle of the fin relative to the horizon to effectively generate lift (Eiffel 1913; Anderson 2005; Fossati 2009), we also quantified incidence angles of the dorsal and anal fins. Additionally, we gathered ecological habitat-usage data from the literature to place locomotor morphology into the context of flow regime differences between ecophases. We then subjected phenotypic data to a series of phylogenetic comparative analyses to assess differences in the tempo and mode of phenotypic diversification between the two ontogenetic stages. Our study provides a detailed investigation of the impact of life stage on the phenotypic diversification within a lineage of fishes, giving a critical perspective on the ontogenetic decoupling of macroevolutionary dynamics in the evolutionary radiation of marine biodiversity.

## Methods

### Data Acquisition

#### Compiling digital images of specimens and habitat usage patterns

Images of 270 adult triggerfishes spanning 26 species in all major triggerfish lineages were from Dornburg et al. (2011). An additional set of 454 images of juvenile and adult fishes were taken from online photo repositories and digitized specimens from the North Carolina Museum of Natural Sciences (**Supplemental Table 1**). This combined dataset encompasses 724 images from 32 triggerfish species (*i.e.* 78% of the extant diversity of Balistidae; **Supplemental Table 2**). All museum specimens were photographed facing left, using 12 megapixel DSLR cameras with a macro lens and ring flash. Juvenile triggerfishes possess unique morphologies and spot patterns still visible in museum specimens allowing for identification (Lyczkowski-Shultz et al. 2003). Habitat data for adult and juvenile life stages were compiled from the literature (**Supplemental Table 2**). Habitats used by triggerfish during their adult and juvenile life stages encompass the majority of tropical and temperate marine habitats (**Supplemental Table 2**). To characterize this broad range of habitats in a manner meaningful to locomotor function, habitat occupancy was partitioned into three bins: (1) mangroves, lagoons, and other structurally complex 3D matrices that buffer against pulses of high water flow; (2) open habitats such as surge zones, outer reefs, or the off-shore pelagic realm; and (3) both (1) buffered and (2) high flow/open habitats.

#### Quantifying Locomotor Morphology

Landmark-based geometric morphometric methods were used for shape quantification (Zelditch et al. 2012a). Landmark placement on fish bodies and fins was accomplished in the TpsDIG2 software **(Rohlf 2006)**. To quantify body shape, 23 homologous landmarks also used by Dornburg et al. (2011) as well as other studies of acanthomorph body shape diversity (Aguilar-Medrano et al. 2016; Collins et al. 2016) were used to capture body shape variation (**Supplemental Fig. 1A**). These are: (1) the posteroventral corner of the maxilla; (2) the anteroventral tip of the premaxilla; (3) the anterodorsal point of mouth where the fleshy lip meets scales; (4) the most anterior point of the eye; (5) the most dorsal point of the eye; (6) the most posterior point of the eye; (7) the most ventral point of the eye; (8) the center of the eye; (9) the anterior point of the first dorsal spine insertion; (10) the dorsal fin origin; (11) the anterior point where the dorsal fin sheath joins the fin rays; (12) the dorsal fin insertion; (13) the dorsal inflection of the caudal peduncle; (14) the dorsal caudal fin ray insertion; (15) the ventral caudal-fin ray insertion; (16) the ventral inflection of the caudal peduncle; (17) the anal fin insertion; (18) the anal fin origin, (19) the posterior inflection where the pelvic flap meets the body; (20) the posterior point of the pelvic spine insertion; (21) the anterior point of the pelvic spine insertion; (22) the posterior ventral point where the fleshy lower lip meets the scales; (23) the anterodorsal point of the lower jaw. To better capture the curves of the body between landmarks, the five sliding semi-landmarks were placed as follows: (1) at the midpoint between landmarks 3 and 9; (2) at the midpoint of the dorsal fin (placed along the body); (3) at the midpoint between the dorsal and ventral caudal fin ray insertions (placed along the fin ray insertion margin); (4) at midpoint of the anal fin (placed along the body); and (5) at the midpoint between landmarks 21 and 22.

For the dorsal and anal fins, a total of four landmarks were placed on each image: insertion and tip of the anterior soft fin ray, insertion and tip of the posterior soft fin ray. Sliding semi-landmarks were placed to capture the curvature of each fin’s distal margin. These were placed by resampling a curve drawn to an outline of each fin’s distal margin divided into nine sliding semi-landmarks (Bookstein 1997). In addition, a sliding semi-landmark was placed along the fin base at the midpoint between the fin origin and posterior insertion for a total of 13 fin landmarks (four fixed, nine sliding semi-landmarks; **Supplemental Fig. 1B**). Caudal fins could not be photographed consistently due to a high frequency of damage in the specimens and we thus restricted analyses to dorsal and anal fins. Since these two median fins represent the primary axes of locomotion in triggerfishes (Sfakiotakis et al. 1999), omission of the caudal fins is unlikely to represent a major gap in modeling the ecomorphology of these fishes. We calculated the angle of the dorsal and anal fin of each specimen relative to the horizon using the angle formed at the junction of two lines: (1) a line drawn from the anterior fin insertion point to the posterior fin insertion point, and (2) a line drawn parallel to the horizontal plane from the posterior fin insertion point to the anterior region of a specimen (**Supplemental Fig. 1B**). The aspect ratio of each fin was defined following Walker and Westneat (2002) as two times the leading edge of the fin squared, divided by the total fin area (**Supplemental Fig. 1B**). All angles and aspect ratios were measured using the ImageJ software package (Abramoff 2007; Hartig 2013).

### Divergence time estimation

We assembled a phylogenetic dataset by combining publicly available DNA sequence data (Supplemental Table 1) from all previous investigations of triggerfish evolutionary relationships (Holcroft 2004; Dornburg et al. 2008, 2011; Santini et al. 2013; McCord and Westneat 2016b). This dataset included DNA sequence data from 11 genes (12S, 16S, RAG1, RAG2, rhod, Tmo-4c4, Bmp4, COI, CytB, Glyt, MYH6, RAG2) for a total of 9,108 base pairs (bp). 12S and 16S were aligned to structural models following Dornburg et al. (2008). Multiple sequence alignments for all other loci were conducted using MAFFT (Katoh et al. 2009) in Geneious v.7.1.2 with alignments confirmed by eye. The combined dataset included a total of 36 triggerfish species that span all major lineages as well as 46 monacanthid, two molid, and one diodontid species (**Supplemental Table 3**) that facilitated the use of fossil calibrations also used by Santini et al. (2013).

Divergence times and phylogenetic relationships were simultaneously estimated in a Bayesian framework using BEAST v2.4.5 (Cummings 2004; Drummond and Bouckaert 2015). Two independent analyses were run for 50 million generations logging parameter estimates every 1000 generations. Analyses were run under an uncorrelated lognormal relaxed clock with a birth-death speciation prior on branching times. Substitution and clock models were unlinked among genes and between stem and loop regions of the ribosomal genes with each partition assigned a GTR + I + Γ_4_ model of sequence evolution. Convergence between runs and appropriate burn-in levels were assessed through visual inspection of the log-likelihoods and parameter estimates in Tracer v1.7 (Rambaut et al. 2018). For each run, effective sample size (ESS) values were quantified to assess effective sampling of the posterior distribution [ESS values > 200 were taken as indicators of adequate sampling (Drummond et al. 2006). As data partitioning can also have a major impact on phylogenetic inference (Kainer and Lanfear 2015), we partitioned our alignment using Akaike Information Criterion (AIC) in PartitionFinder2 using the greedy algorithm (Lanfear et al. 2017). The candidate pool of possible partitions contained each gene’s separate codon positions as well as the stem and loop regions of each ribosomal gene (**Supplemental Table 4**). Substitution and clock models were unlinked following selection of the best-fit partition model.

To time-calibrate the phylogeny, we followed the fossil calibration strategy used by Santini et al. (2013). Briefly, this approach used the stem balistid *Gornylistes prodigiosus* to calibrate the divergence between triggerfishes and filefishes. This fossil from the Northern Caucasas is Middle Eocene in age (41-42 Ma) and represents the oldest known crown balistoid (triggerfish+filefish) fossil (Bannikov and Tyler 2008). The 95% soft upper bound on the prior age calibration for this split was determined using the presence of the stem Balistoid *Bolcabalistes* at 50 Ma (Santini and Tyler 2004). *Austromola (Gregorova et al. 2009)* was used to calibrate the split between *Mola* and *Ranzania* at 22 Ma, with a soft upper bound based on *Eomola* (Tyler and Santini 2002). A minimum root age for the tree was set to 59 Ma based on the appearance of the Danish fossil *Moclaybalistes danekrus* with a soft upper bound of 85 Ma based on the appearance of *Protriacanthus gortani* (Tyler and Santini 2002).

Santini et al. (2013) suggested that more inclusive sampling of monacanthid taxa improved site rate estimates and explains the inconsistency of divergence times between studies of triggerfishes. However, nucleotide substitution saturation could provide an alternate explanation. To test the impact of saturation on our branch length estimates, we quantified site rates of each locus using Hyphy (Pond et al. 2004) in the PhyDesign web interface (López-Giráldez and Townsend 2011) using the time tree from Santini et al. (2013) as a guide. Using the estimated site rates, phylogenetic informativeness (PI) profiles were calculated using the R package PhyInformR (Dornburg et al. 2016). Declines in the shape of PI profiles have been found to reflect intervals where branch length estimates have become compromised (Dornburg et al. 2014, 2017b). As such, we profiled each codon position of each gene as well as the stem and loop regions of each ribosomal gene, excluding loci from selected partitions that possessed a sharp decline in informativeness prior to the root of the guidetree (Dornburg et al. 2014). It should be noted that informativeness calculations have been found to be robust between input guidetrees that vary in topology or branch lengths (Dornburg et al. 2017a), so the use of this guidetree over alternative divergence time estimates would be expected to produce indistinguishable PI profile shapes. The resulting alignment was then subjected to two BEAST analyses that mirrored the conditions outlined above. All analyses were also repeated using the same BEAST conditions and filtration criteria as above on an unpartitioned alignment to assess the additional possible effect of partition choice on branch lengths.

### Variation in the diversification of form and function between life stages

#### Life stage and morphospace occupation

For each species and shape dataset, a Procrustes fit was used to initially remove variation due to scaling, rotation, and translation in the landmark based data (Rohlf and Slice 1990; Zelditch et al. 2012a), generating mean shape datasets for each species at each life stage (juvenile or adult). The mean configurations of each species were subsequently combined, and subjected to a second Procrustes fit and relative warps analysis in the TPS relwarp software (Rohlf 2007). To visualize patterns of trait diversity for all data and to qualitatively compare them between the two ontogenetic stages, phenograms were plotted for each trait in both adult and juvenile stages using the R-package phytools (Revell 2011). A distance-based Procrustes ANOVA (function *procD.lm*; 9999 iterations with a randomization of raw data) available in the R-package geomorph (Adams and Otárola-Castillo 2013) identified the significance in divergences between adults and juveniles in trait spaces. We then assessed phylogenetic signal to compare the pattern of morphological evolution between ecophases. We used the generalised K statistic, also implemented in the R-package geomorph (Adams and Otárola-Castillo 2013), that can be applied on univariate and multivariate data (Adams 2014a). A *K* value greater than 1 implies that closely related species are more similar than expected under a null model of Brownian motion, whereas a value that is less than 1 suggests that relatives resemble each other less than expected. Rather than absolute values of *K* on this scale, we focus on how *K* differs between juveniles and adults. Statistical significance was assessed by permutation (1,000 iterations).

#### Life stage and disparity

No evidence has been presented for a linear relationship between the pattern of morphological disparity and ontogeny in teleost fishes (Zelditch et al. 2003; Frédérich and Vandewalle 2011). Thus we might also expect an absence of relationship between juvenile and adult morphological disparity in triggerfishes. To test this hypothesis, we first calculated the level of shape disparity based on Procrustes variance (Zelditch et al. 2012b) and performed comparisons between ontogenetic stages (i.e. permutation test, 9999 iterations) using the function *morphol.disparity* in the R package geomorph (Adams and Otárola-Castillo 2013).

Next we quantified and contrasted calculations of relative subclade disparity through time (DTT Harmon et al. 2003) for both stages. Previous work suggested that between-clade disparity arose early in the evolutionary history of triggerfishes as a consequence of divergent fin aspect ratios allowing early lineages to exploit a range of marine environments (Dornburg et al. 2011). However, it is not clear if patterns of morphological disparity between juveniles follow a similar pattern. Accordingly, we compared whether juvenile fin aspect ratios depict a similar pattern to adult disparity by calculating the relative subclade disparity through time (Harmon et al. 2003) for both stages. We compared results to the expectations of a null model of Brownian motion generated from 10,000 simulations (Slater et al. 2010). We additionally conducted analyses of DTT on fin insertion angles relative to the horizon to test the expectation that this aspect of body shape should mirror patterns of disparity observed in aspect ratios. As even for traits evolving under pure Brownian motion, principle components and similar ordination analyses such as a relative warps analysis can bias the major axes towards displaying a signature of an early burst of trait evolution (Uyeda et al. 2015), additional shape data was not used in the DTT analyses. All analyses were conducted in the R package geiger2 (Pennell et al. 2014).

In addition to quantifying patterns of subclade disparity through time, we also used the approach from Cooney et al.(2017) to quantify global patterns of morphospace occupancy of aspect ratios and incidence angles through time. Maximum-likelihood based ancestral states for each trait in either the juvenile or adult dataset were first estimated using the R package phytools (Revell 2011). Ancestral disparity through time values were based on one million year time slices and calculated as the sum of the variances for each trait in each time slice. Empirical disparity values were compared to trait values generated under a null Brownian model of evolution. Differences in the Loess-fitted slope of disparity between the null model and each dataset were then computed using the R package msir (Scrucca 2011). A difference of zero indicates no departure from the expectation of Brownian motion, while a positive difference indicates an increase in morphospace expansion and a negative difference indicates the packing of existing morphospace (Cooney et al. 2017). For each trait, adults and juveniles were compared to assess differences in morphospace occupancy between life stages.

#### Life stage and rates of trait evolution

Varied natural selection operating over ontogeny can promote variation in the dynamics of morphological evolution between life stages. We tested this prediction by first comparing the overall net rates of morphological diversification between adults and juveniles with the function *compare.multi.evol.rates* (Adams 2014b) from the R-package geomorph using a Brownian motion model of trait evolution. This approach calculates the net rate of morphological evolution (σ^2^) for each trait from the data, allowing a ratio of rates to be quantified, with significant differences assessed via phylogenetic simulation (10,000). The comparisons of overall net rates were performed for every morphofunctional trait, including univariate and multivariate data. For comparing the evolutionary rates of multivariate traits, we followed the procedure of Denton and Adams (Denton and Adams 2015).

In addition, we conducted a series of Bayesian analyses of macroevolutionary mixtures (BAMM) (Rabosky 2014) to compare rates of phenotypic evolution between ecophases. For each data type (adult versus juvenile) and trait dataset (fin aspect ratio and incidence angle), we set prior parameters using the R package BAMMtools (Rabosky et al. 2014). All analyses were run twice independently for 50 million generations, sampling every 1,000 generations. Convergence between runs was assessed through visual inspection of the log-likelihoods and effective sampling of the target posterior distribution of parameter values was assessed through quantification of ESS values (ESS >200). Finally, we used tip-specific BAMM estimates of trait evolutionary rates, which are the mean of the marginal posterior distribution of rates for individual species. We conducted an ordinary least squares regression analysis on tip rate values to test the relationship between the dynamics of morphological evolution observed at both stages. In all cases, BAMM analyses were conditioned on taxon sampling at both adult and juvenile stages. Phenotypic data were available for a larger set of species at the adult stage than for juveniles (**Supplemental Table 5**). To assess the sensitivity of our results to uneven taxon sampling levels between juveniles and adults, analyses were repeated following two schemas: (1) we assessed phenotypic rates for individual species when using taxa shared by adult and juvenile datasets and we then performed the tests of linear relationships; (2) we quantified phenotypic rates for individual species when including the maximum of taxa for both adult and juvenile datasets, then we dropped species for which morphofunctional data are not available at both ontogenetic stages and we finally performed ordinary least squares regression analysis.

## Results

### Estimates of triggerfish divergence times and patterns of habitat use between ecophases

Across all analyses, the estimated phylogenetic tree topology mirrors previous studies of phylogenetic relationships by supporting six major clades: (1) *Abalistes*; (2) *Canthidermis*; (3) *Sufflamen*; (4) *Rhinecanthus*; (5) *Balistes*; and (6) all other balistids (Dornburg et al. 2008, 2011; Santini et al. 2013; McCord and Westneat 2016b). Using all possible subsets of data under a best-fit partitioning strategy (**Supplemental Table 4**), posterior probability values for all but one node (placement of *Rhinecanthus verrocosus*) was greater than 0.95 (**Fig. 1, Supplemental Fig. 2**). PI profiles of several data partitions reached their apex prior to the root of the guide tree (**Supplemental Fig. 2B**). A decline in PI following the apex of a profile has been dubbed a “rainshadow of noise”, the signature of an increase in the number of hidden substitutions (Townsend and Leuenberger 2011; Dornburg et al. 2017b) that can mislead branch length estimation (Dornburg et al. 2014). Exclusion of data that depicted declines in PI profiles resulted in very different, and at times mutually exclusive, highest posterior density intervals (HPD) of node ages (**Supplemental Fig. 2**). In general, posterior age estimates were older, generally on the order of 4-5 Ma, when all data was included. The HPD interval of the estimated age for the most recent common ancestor (MRCA) of crown triggerfishes ranged between 21.3 and 24.6 Ma (mean: 22.9 Ma) when all data was included and between 15.9 and 9.4 (mean: 12.6) when data was partitioned and filtered (**Fig. 1 and Supplemental Fig. 2**).

**Figure 1:**
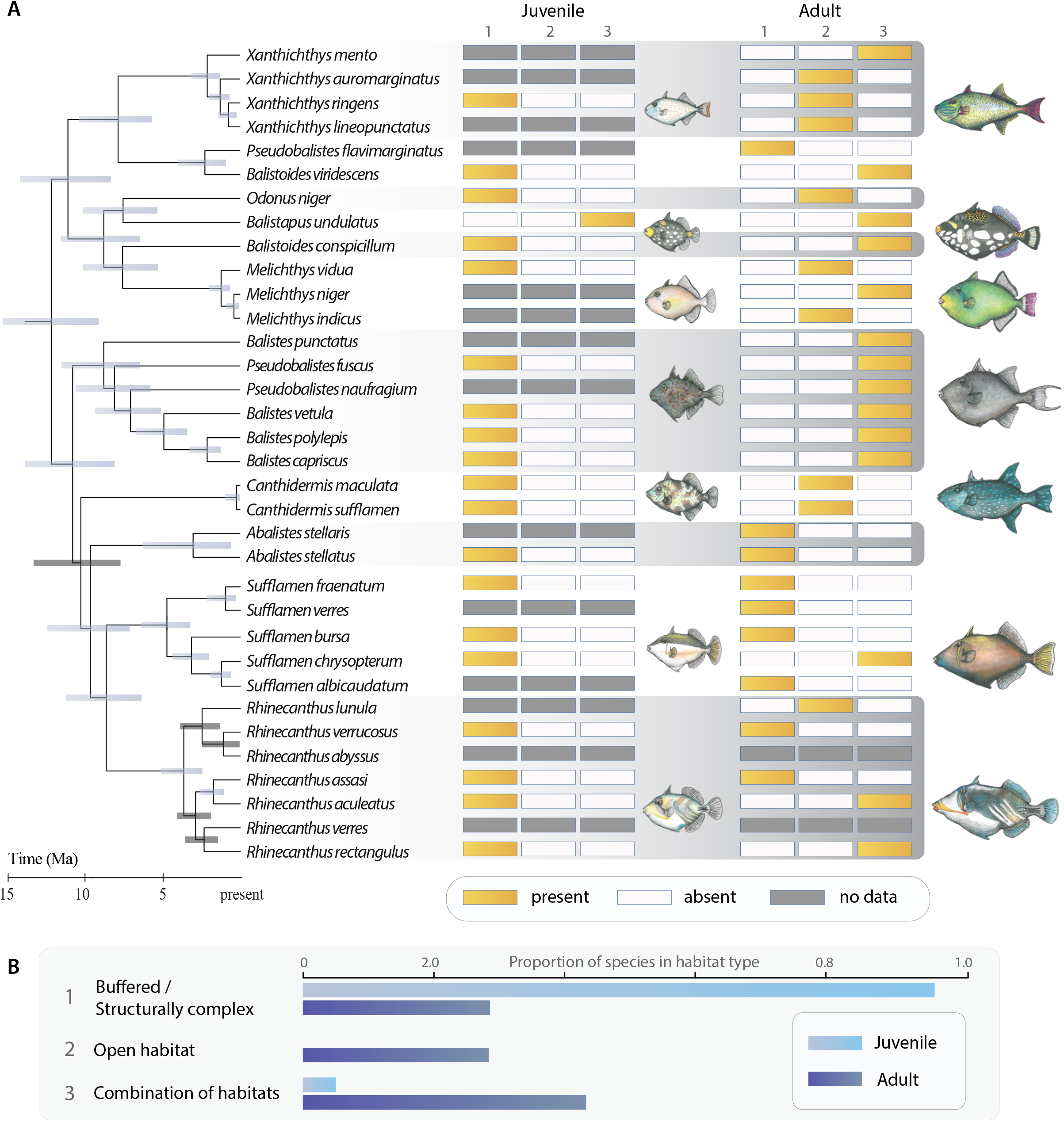
A phylogenetic perspective of triggerfish habitat use between life stages. A. Divergence time estimates of Balistidae from phylogenetically informative partitions and visualization of habitat types between life stages. B. Proportion of species per habitat type between life stages. Numbers in A (1,2, or 3) correspond to habitats in B. Illustrations by K. Zapfe.

We found that juveniles from nearly all lineages were associated with low flow habitats. However, our results reveal a history of independent colonization of open water habitats across divergent triggerfish lineages. Placing ecological data into this evolutionary context revealed repeated convergences into habitats characterized by high flow or a combination of high and low flow in adult triggerfishes (**Fig. 1**). For example, the open habitat was invaded by different lineages: *Xanthichthys*, *Canthidermis*, *Melichthys* and *Odonus* (**Fig. 1A**). *Rhinecanthus* showed the largest diversity of habitat usage at the adult stage, having representative lineages restricted to more open versus structurally complex habitats as well as lineages that used a combination of the two categories (**Fig. 1**).

#### Life Stage and the diversification of locomotor morphology

Visualizations of phenograms revealed major differences in overall patterns of morphospace occupancy between juvenile and adult triggerfishes (**Fig. 2**). The functional morphology of triggerfishes varied significantly between the two life stages (**Table 1**). Juveniles, which were found almost exclusively in sheltered habitats subject to low flow regimes, only possess low aspect ratio fins and thus occupy a fraction of the aspect ratio diversity found in adults (**Fig. 2A and 2B, Table 2**). Correspondingly, dorsal and anal fin shape was primarily captured in changes in the length of the fin rays that formed the leading edge of each fin on the first RW that explained 49.8 % and 38.2 % of the total variation between adult and juvenile dorsal and anal fin shapes respectively (**Fig. 2C and 2D; Supplemental Table 5**). The second major axis of dorsal and anal fin shape variation (RW2) explained change in curvature of the distal fin margins and respectively accounted for 24.3% and 35.2% of the variation (**Supplemental Table 5**). Qualitatively, variation along RW1 and RW2 axes is suggestive of a higher level of shape diversity at the adult stage (**Fig. 2C and 2D)**. However, when considering all shape information, this difference in disparity levels between life stages was not significant (**Table 2**).

**Figure 2.**
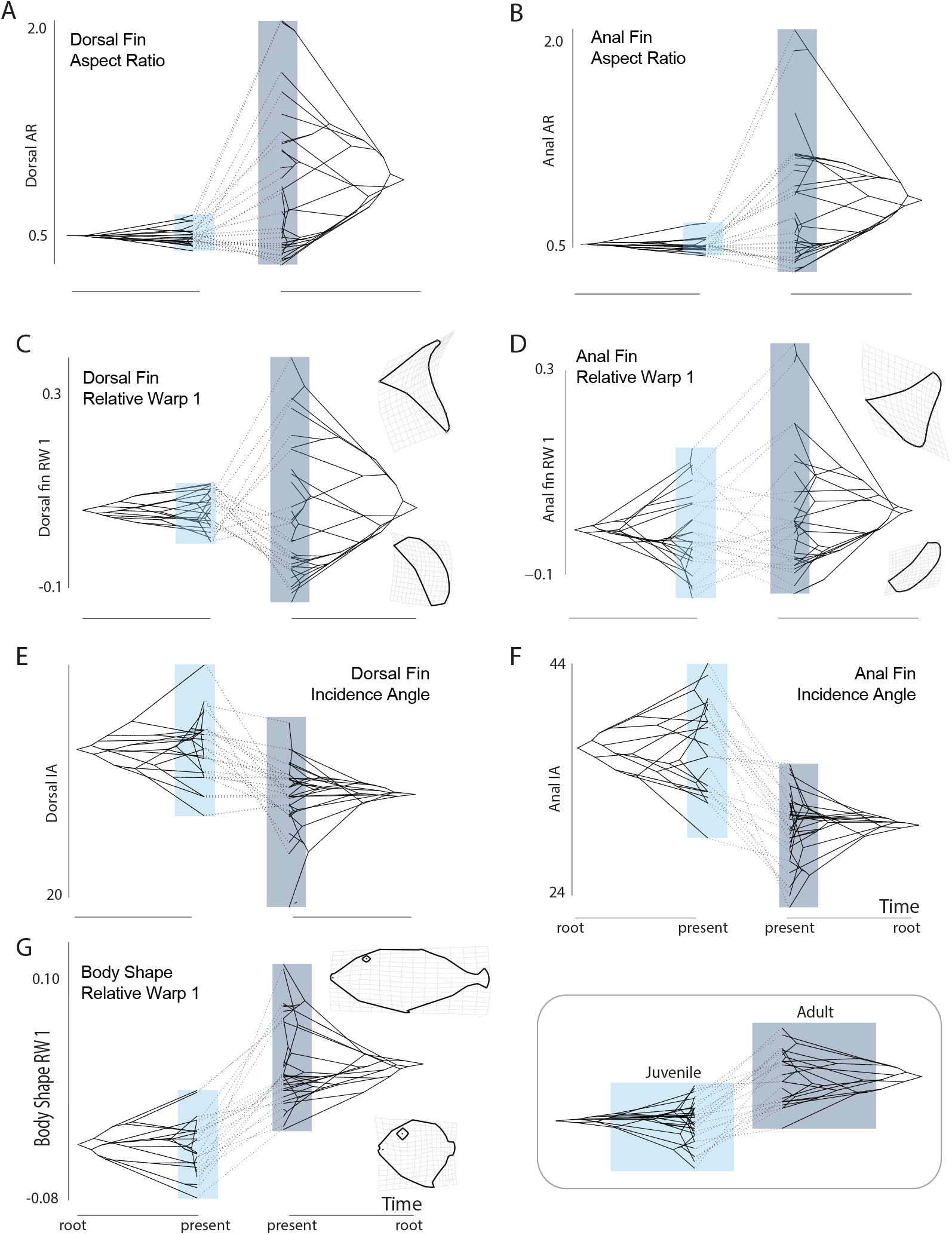
Phenograms comparing morphospace occupancy between juvenile (left each panel) and adult (right in each panel) triggerfishes for (A) dorsal fin aspect ratio; (B) anal fin aspect ratio; (C) the first relative warp axis of dorsal fin shape variation; (D) the first relative warp axis of anal fin shape variation; (E) Dorsal fin incidence angle; (F) anal fin incidence angle; and (G) the first relative warp axis of body shape variation. Light blue boxes indicate the range of trait values for juveniles, dark gray the range of adults in each panel.

In contrast to fin shape and aspect ratios, the scope of diversity of incidence angles in juvenile triggerfishes is on par with that observed in adults, albeit shifted towards lower angles (**Fig. 2E and 2F)**. These changes in angle correspond to overall changes in body shape between adults and juveniles. A dorsal-ventral compression from a discoid juvenile shape to more fusiform adult body shape is observed, which is coupled with an elongation of the anterior cranium forming the primary axis of body shape change accounting for 41.2 % of the variation (**Fig. 2E**). The second major axis of body shape variation (20.0 % of the total shape variation) is primarily associated with an elongation of the head region (**Supplemental Table 3**).

This qualitative description of variation in the pattern of diversification through trait spaces between stages is strengthened by the tests of phylogenetic signal (**Table 3**), that demonstrate opposing evolutionary patterns between ecophases for fin aspect ratios. In adults, closely related species are more similar than expected by chance, while in juveniles these traits are more evolutionary labile (Dorsal ratio: K__adult__ = 1.2 Vs K__juvenile__ = 0.6; Anal ratio: K__adult__ = 1.6 Vs K__juvenile__ = 0.9). Similarly, quantification of phylogenetic signal for body and anal fin shapes was low in juveniles and moderate in adults (**Table 3**).

We found that adults possess a higher level of disparity in fin aspect ratios than juveniles (p < 0.01, Fig. 2), but no significant differences between ecophases were observed for the other traits (**Table 2**). Our results quantifying overall patterns of disparity over time revealed that dorsal and anal aspect ratios in both adults and juveniles were generally similar. In both cases, disparity was near the expectations under Brownian motion, packing existing morphospace following the early diversification of the crown group (**Fig 3A & 3B**). In contrast, disparity in juvenile incidence angles reflects an expansion of morphospace over time (**Fig 3C & 2D**) that sharply diverges from the pattern of less morphospace occupancy over time than that expected under a Brownian model observed for adults (**Fig 3C and 3D**).

**Figure 3.**
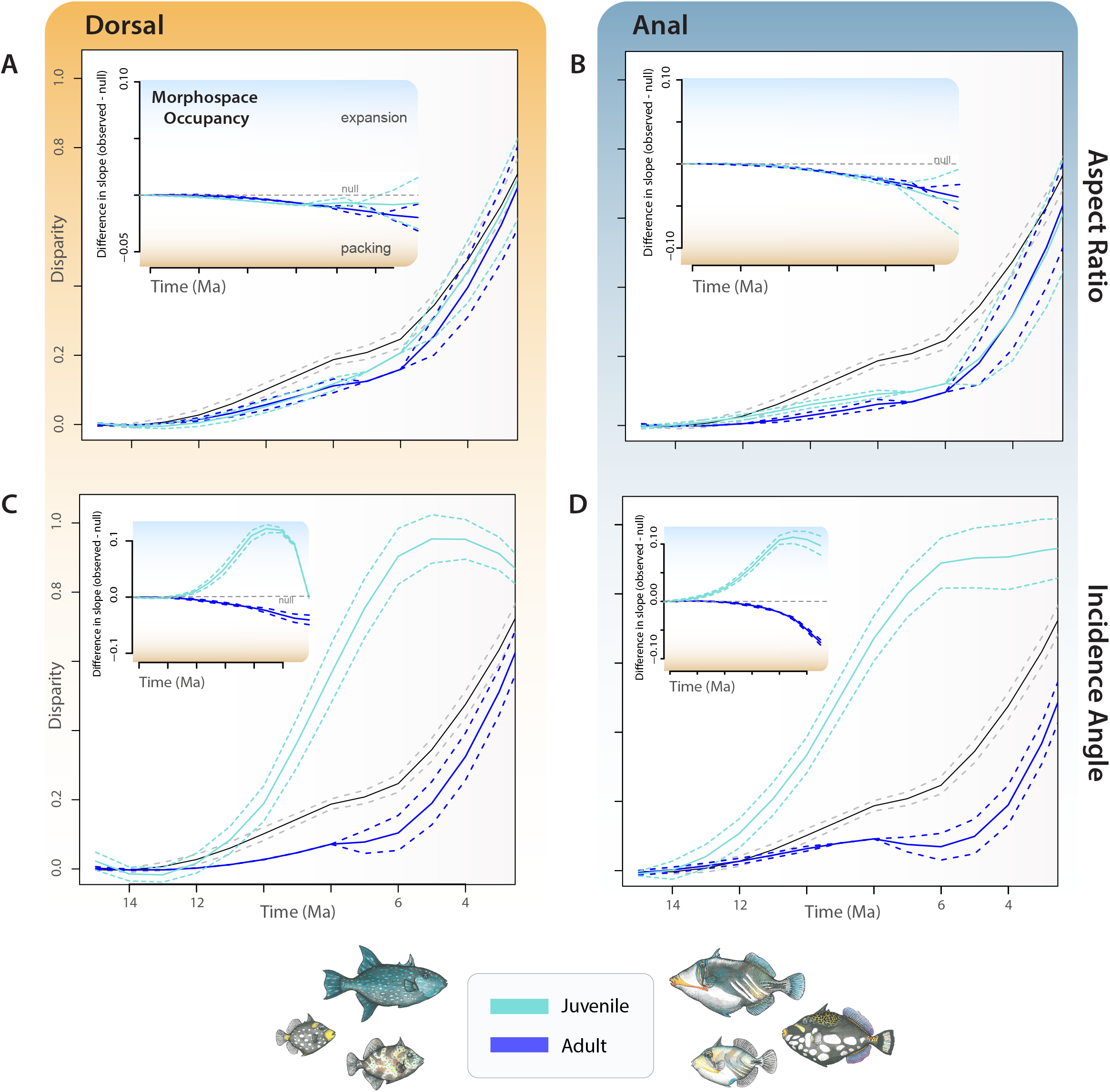
Contrasting patterns of the accumulation of disparity and morphospace occupancy (insets) through time between adults (dark blue), juveniles (aqua), and a null expectation of disparity based on brownian motion (gray) for (A) dorsal fin aspect ratio; (B) anal fin aspect ratio; (C) dorsal fin incidence angle; and (D) anal fin incidence angle. For insets, positive differences in slope indicate expansion of trait diversity (light shading) while negative differences indicate “packing” of an already occupied region of morphospace (orange shading).

Placing these findings into the context of subclade disparity mirrors previous investigations of locomotor disparity in adults for aspect ratios (Dornburg et al. 2011): disparity is partitioned between subclades early in the evolutionary history of the clade (**Fig 4A & 4B**). A similar pattern is observed for the anal fin aspect ratio in juveniles (**Fig. 4B**), while subclade disparity within clades is higher for the dorsal fin aspect ratio (**Fig. 4A**). In contrast, subclade disparity patterns of incidence angle revealed higher than expected disparity within subclades that exceeded expectations of the Brownian null model (**Fig 4C & 4D**). In addition to divergent patterns of morphospace occupancy, the dynamics of morphological evolution are also highly discordant between juvenile and adult stages. Except for the incidence angles, the overall net rates of evolution significantly differ between stages (**Table 4**). The rate parameters (σ^2^) are higher in adults in comparison with juveniles for the dorsal fin shape and both fin aspect ratios, with juveniles exhibiting higher rates of morphological diversification for the body and anal fin shapes than adults (**Table 4**).

**Figure 4.**
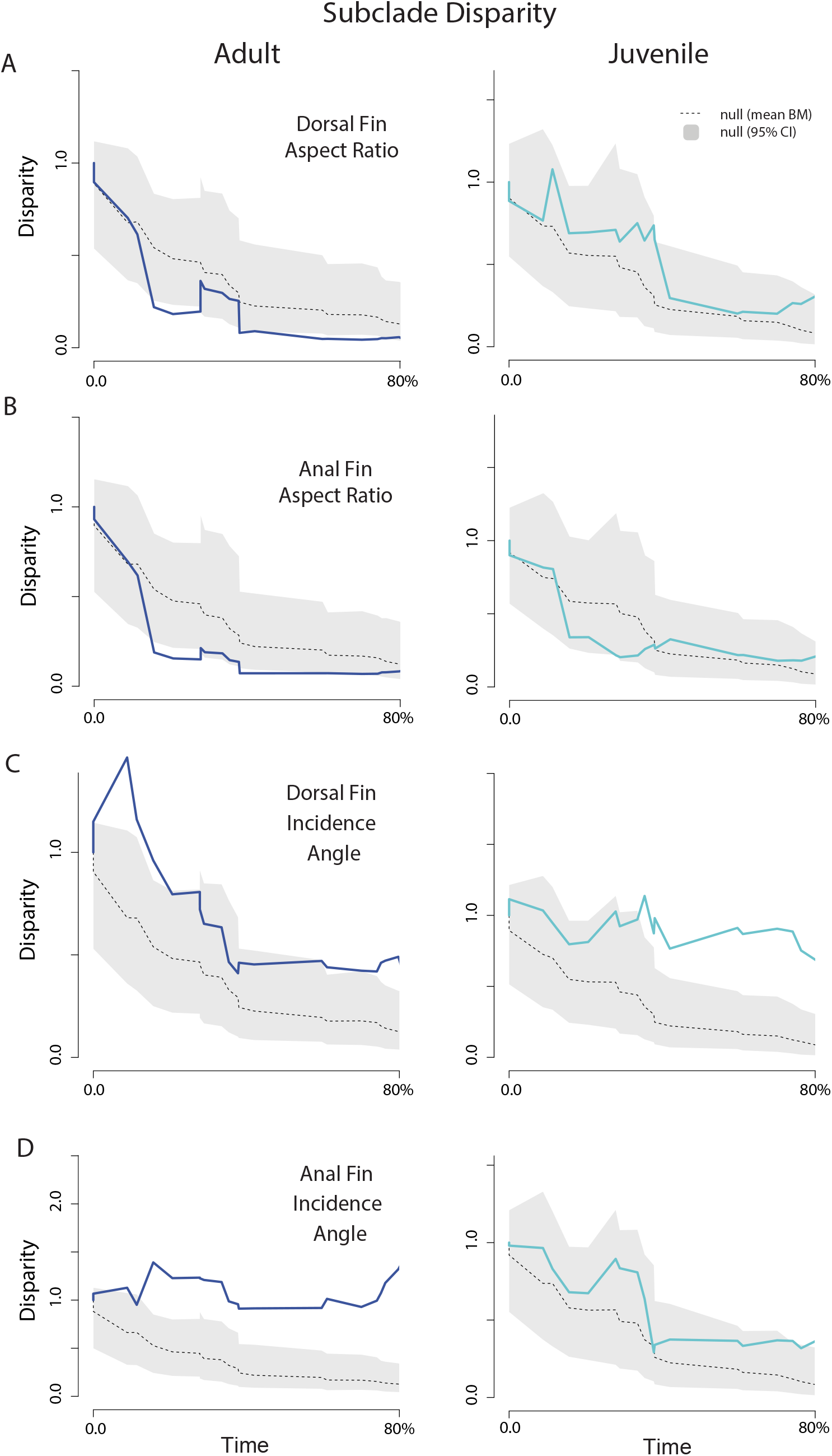
Subclade disparity through time for adults (dark blue), juveniles (aqua), and a null expectation of disparity based on brownian motion (gray) for (A) dorsal fin aspect ratio; (B) anal fin aspect ratio; (C) dorsal fin incidence angle; and (D) anal fin incidence angle. Dotted gray line indicates the mean subclade disparity through time across the brownian motion simulations.

Results from BAMM further demonstrate a decoupling of macroevolutionary dynamics between adults and juveniles and between rates of aspect ratio and incidence angle diversification (**Fig. 5**). The relationships between the estimated rates of morphological evolution in juveniles and adults are poor or even non existent. All linear models were not significant, except for dorsal fin aspect ratios when rate estimates are calculated with the maximum taxon sampling for both stages (**Table 5**). A visual exploration of phylorates revealed that dorsal and anal fin aspect ratio diversification is similar in adults, with higher rates in the early history of the group (**Fig. 5A & 5B**). In contrast, juvenile rates are fairly uniform and over an order of magnitude slower than those estimated for adults (**Fig. 5A & 5B**). Differences in the rates of change for incidence angles between juveniles and adults are similar (**Fig 5C & 5D**). Again rates are faster in the early history of the group for adults at levels that vastly exceed the rates estimated in juveniles. Additionally, there is a signature of an acceleration of rates towards the present diversification in the adult incidence angle phenotype of the primary open habitat associated lineage *Canthidermis*. In all cases, these results were robust to taxon sampling strategy (**Supplemental Fig. 3 and Table 5**).

**Figure 5.**
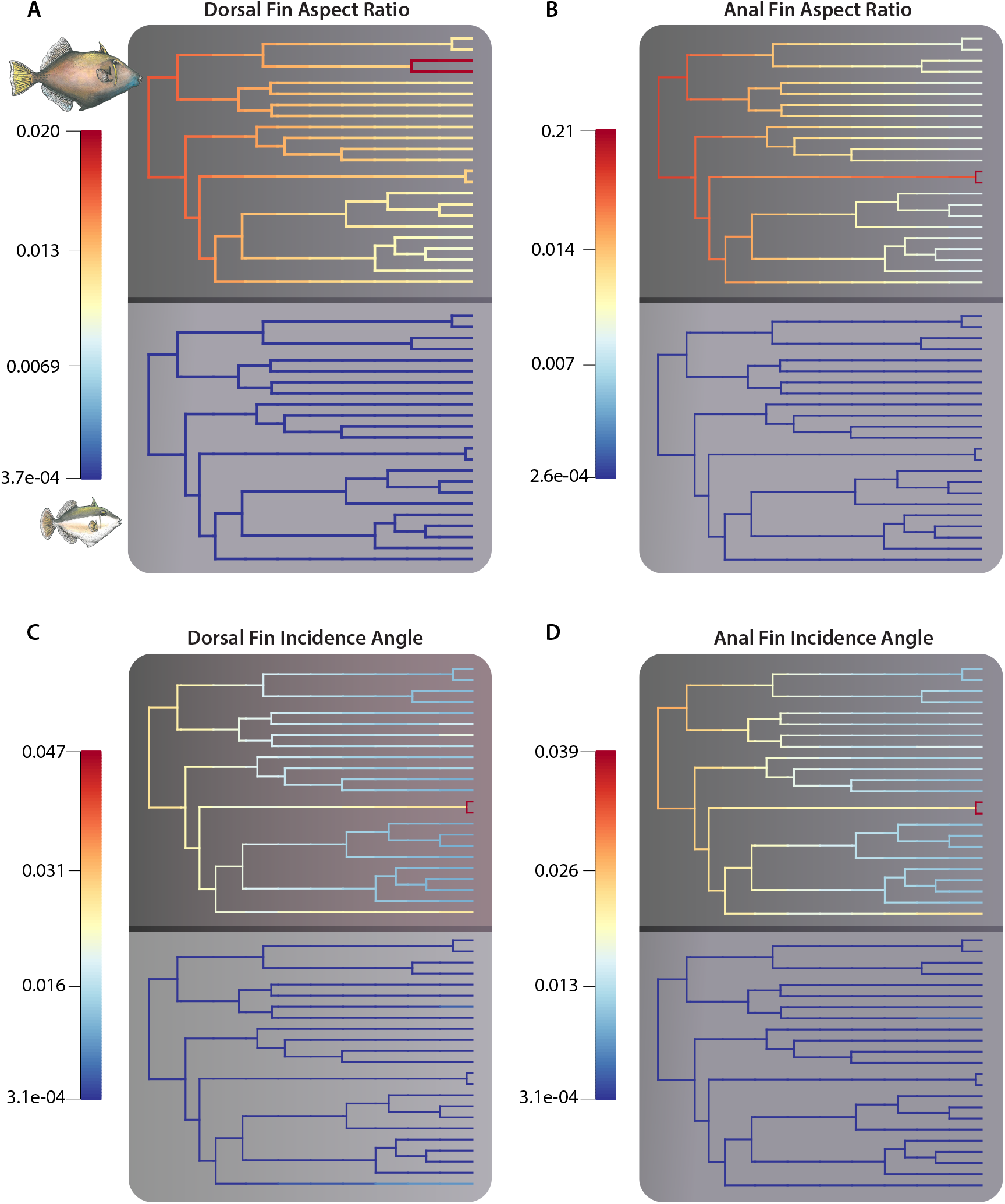
Visualization of rates of trait diversification between adults (top) and juveniles (bottom) for (A) dorsal fin aspect ratio; (B) anal fin aspect ratio; (C) dorsal fin incidence angle; and (D) anal fin incidence angle.

## Discussion

Much of our understanding of the rules that govern the origin and maintenance of phenotypic diversity in marine fishes has been based on macroevolutionary studies focused on only adults. However, all of our trait-based analyses demonstrate unique patterns of phenotypic diversification between adult and juvenile life stages of triggerfishes. The results of our literature search reveals that juveniles are more restricted in their habitat use than adults, corresponding to a striking difference in fin and body shape morphologies between the two life stages. For adult phenotypes, this difference in morphology reflects an early divergence of fin and body shapes between major lineages. Such a difference is not mirrored at the juvenile stage. Instead, juvenile forms either have a restricted morphospace or consistently expand in morphospace over time with closely related species that are often highly disparate. Correspondingly, the rate of phenotypic diversification for juveniles is generally an order of magnitude lower than that estimated for adults. Collectively, these results illustrate how heterogeneity in life history can decouple the diversification dynamics between ecophases in coastal fishes.

### Life Stage and Disparity of Triggerfish Locomotor Morphology

The results of every morphospace and disparity analysis strongly support distinct patterns of disparity between adult and juvenile triggerfishes. Juveniles lack high aspect ratio fins required for prolonged cruising and generally possess more discoid body shapes (**Fig. 2**). Low aspect ratio fins are associated with increased maneuverability required in 3D nursery environments (Fulton et al. 2005) and our visualizations of aspect ratio evolution through time demonstrate a pattern where juveniles are restricted to a small range of low aspect ratios (**Fig. 2A & 2B**). However, juvenile fin incidence angles varied between species substantially with juveniles expanding their diversity of incidence angles over time, often as a result of divergence between closely related species (**Fig. 2E & 2F**). In general adults possess lower incidence angles than juveniles as a consequence of having higher aspect ratio fins, and this shift in incidence angles between juveniles and adults is also reflected in body shape changes. Juveniles generally possessed more discoid body shapes while adult stages exhibited more fusiform shapes (**Fig. 2**).

Several hypotheses can be invoked to explain these changes in locomotor morphology between ecophases. First, it is possible that multiple combinations of body shapes and fin incidence angles could produce similar locomotor performance with a given low fin aspect ratio. Such a many-to-one mapping between form and locomotor performance would allow for the evolution of a diverse array of phenotypes (Alfaro et al. 2005; Wainwright et al. 2005; Zelditch et al. 2017). Recent studies have demonstrated that for aspect ratios greater than 0.75, incidence angles above 35° tend to disrupt flow and therefore the ability to generate lift (DeVoria and Mohseni 2017). While aspect ratios of 0.75 are certainly low from an engineering standpoint, we found the median aspect ratio for juvenile triggerfishes across all species to be 0.49, with half of all sampled taxa possessing aspect ratios below 0.50 (**Fig. 2**). Additionally, airfoils are often studied in isolation or assumed symmetric in engineering studies (DeVoria and Mohseni 2017). However, this is not the case in triggerfishes. Although fin evolution is correlated between the median fins (Dornburg et al. 2011), high aspect ratio swimmers tend to have more pronounced dorsal fins compared to anal fins (**Fig. 2**), and anal fin shape tends to be more variable than dorsal fin shape in juveniles (**Fig. 2, Table 2**). Continued theoretical studies, swimming performance studies, and hydrodynamic modeling of asymmetric and low aspect ratio fins in the range of those observed in triggerfishes are required to evaluate how incidence affects performance and whether body shape is more free to vary among juvenile fishes.

It is also possible that body size limits the ability of juvenile triggerfish to explore open habitats. Small fishes in open marine habitats tend to exhibit specific behaviours while schooling to avoid predation (Magurran 1990). Triggerfishes are not known to exhibit schooling behaviours, which could render juveniles highly vulnerable to predation in open habitats. An alternative, but not mutually exclusive hypothesis to both predation, as well as a many-to-one mapping of form and function, is the possibility that complex 3D environments might relax selection for optimized horizontal movement. This is certainly plausible for adult triggerfishes such as the open habitat-ranging ocean triggerfish *Canthidermis sufflamen* (*Brito et al.* 1995). In this species, a high aspect ratio locomotor morphology associated with cruising is clearly advantageous over a squat, reef-associated form. However, such extremes in locomotor types do not exist among juveniles in nursery habitats. Instead, we propose that changes in body shape/incidence angle more likely reflect avoidance of competition, predation, or exploration of new niche space as species occupy different nursery types with different prey resources. These hypotheses may in part explain the often large divergences in juvenile body shape morphospace/incidence angles between closely related species (**Fig. 2**). Although the feeding ecology of most juvenile triggerfishes has not been studied, the finding of both intraspecific conflict and changes in prey acquisition strategies between sympatric species (i.e., selective picking in juvenile *Rhinecanthus* versus blowing water onto sand to expose prey in juvenile *Balistoides viridescens*) support this hypothesis (Chen et al. 2001). Given that major changes in available prey resources occur between nurseries as disparate as mangroves, lagoons, and sargassum mats in pelagic environments, further fine-scale studies of juvenile triggerfishes within and across different nursery habitats are critically needed to assess whether nursery specific ecological opportunities have canalized adult phenotypes.

#### Considering alternate macroevolutionary pattern between ecophases

Recent work in Anurans has suggested that ecophases of organisms with complex life cycles can exhibit decoupled diversification dynamics and varied phenotypic patterning (Roulants et al. 2011; Sherrat et al. 2017; Valero et al. 2017). Our study provides evidence that this is also the case for juvenile and adult triggerfishes. However, these findings also raise the question of what mechanisms can give rise to such differences. As illustrated by Darwin, juveniles “might easily be rendered by natural selection different to any conceivable extent from their parents”, with differences that are “correlated with successive stages of development” (Darwin 1859; p354). For organisms that undergo discrete ontogenetic shifts, investigating how disparity evolves between these stages represents an exciting frontier for our understanding of phenotypic evolution. We hypothesize that the high level of disparity for fins aspects ratio at the adult stage is related to the large diversity of habitats (**Fig. 1**). It is likely that strong selective forces such as resource availability, competition and predation cause large shifts in the adaptive landscapes between habitats, that in turn promote adaptive evolution of fin and body shapes. Conversely, the morphofunctional diversity of juveniles evolves more slowly with expansions of morphospace between closely related taxa. Such an evolutionary dynamic is in accordance with the expectation that nursery areas provide higher survival rates than offshore habitats due to factors such as high prey density and biomass, low predator abundance, and complex habitat structure (Nagelkerken 2009). As such, the clustering of fin aspect ratios and body shapes of triggerfish juveniles in morphospace may suggest that subtle phenotypic variation or behavioural adaptations would be sufficient to compete in such environments with distinct selective forces from adult habitats. Moreover, these clusters could represent successful morphotypes adapted to specific nursery habitats with low flow regimes, with recent phenotypic divergences among closely related taxa either representing shifts to new habitats or movement to new adaptive zones in a saturated space. Unfortunately, the ecology of juvenile triggerfishes has not been studied in sufficient detail to enable testing such hypotheses at the time of this writing.

In addition to a consideration of ecology, it is important to remember that variation in phenotypic disparity between life stages likely involves numerous other factors that arise at different times both in ontogeny and in phylogeny. As such, patterns of morphological disparity between juveniles and adults need not always be asymmetric in the same direction. For example, work looking at the ontogenetic variation of fish shape phenotypes has resulted in cases with both higher levels of disparity at the juvenile stage (Zelditch et al. 2003) and higher disparity levels at the adult stage (Frédérich and Vandewalle 2011). While lineage specific differences in juvenile and adult disparity may be expected, these findings highlight the research potential of comparative investigations that integrate data from genotypes and phenotypes at different life stages. Such work is already underway, with recent studies highlighting shifts in gene expression between larval, juvenile and adult life phases in cichlid opsins (Carleton et al. 2008), changes in color and types of chromatophores between adult and juvenile dottyback fishes (Cortesi et al. 2016), as well as the impact of epigenetic mechanisms on phenotypic development (Hu and Albertson 2017) to name but a few. Moreover, in Anurans, adult and juvenile phenotypes have not only been found to evolve asynchronously, but exhibit expression of mutually exclusive sets of phenotype-coding genes between life stages (Valero et al. 2017). A similar finding has been documented in cichlids, that further revealed suites of divergent coexpressed genes that were present in early life stages prior to major phenotypic differentiation and that these became more during ontogeny (Fruciano et al. 2019). Given the ubiquity of fishes that undergo shifts from juvenile to adult forms, similar future studies of gene expression profiles or genome-wide association studies between fish life stages would help illuminate genetic mechanisms facilitating the evolution of phenotypic disparity spanning phases across the Tree of Life (Gaither et al. 2018).

## Acknowledgements

We thank G. Hogue and L. Roupe of the North Carolina Museum of Natural Sciences Ichthyology Research Unit for help with aspects specimen preparation for this project, L. Rao for additional assistance with imaging, B. Stuart for equipment loans, and B. Sidlauskas for insightful conversations during the inception of this project.

