## Supplemental materials for "Considering decoupled phenotypic diversification between ontogenetic phases in macroevolution: An example using Triggerfishes (Balistidae)"

*Systematic Biology*

Supplemental Table 1: Location of specimens examined for this study

| Species | Burke Museum | Bishop Museum | Philadelphia Academy of Sciences | LA County Museum | Yale Peabody Museum | Museum of Comparative Zoology | Field Museum | North Carolina Museum of Natural Sciences |
|---|---|---|---|---|---|---|---|---|
| *Abalistes stellaus* | ---- | --- | 117657 (a-b)<br>117656<br>90440 | --- | --- | 27048<br>24320<br>3758 |  |  |
| *Balistapus undulatus* | UW 009104<br>UW 017709 (a-d) | --- | --- | --- | 7168<br>7169<br>8325<br>9816<br>7167<br>9544 | 63172<br>26356 | 4747<br>89708<br>90739<br>91474<br>110921 | |
| *Balistes capriscus* | UW 005164<br>UW 016887 | --- | 102579<br>102745<br>102579 | --- | 14006 | 62199<br>23972<br>11941<br>26654<br>11946<br>24148<br>54378<br>35799<br>23971 | 4883<br>46697<br>46828<br>48897 | 4551; 4763; 5935; 5990; 7864; 9077; 9132; 9510;<br>11471<br>11705<br>13030<br>15117<br>15587<br>17332<br>10159<br>10171;<br>3976 4906;<br>4985 7875;<br>9816<br>23245<br>27138<br>27157<br>27158<br>27234<br>27244<br>28197<br>28230<br>28275<br>30383<br>30485<br>31204 |

45348
31204
45348
48942
48953
48970
48984
48987
49031
49054
49068
49092
49156
49171
49174
49180
49203
49276
49297
49316
49349
49477
49500
49513
49533
49557
49559
49579
49618
49625
49632
49644
49654
49687
49698
49704
49738
49745
49750
49773
49837
49840
49857
49877

| | | | | | | | | |
|---|---|---|---|---|---|---|---|---|
| | | | | | | | | 49911 |
| | | | | | | | | 49968 |
| | | | | | | | | 49982 |
| | | | | | | | | 49988 |
| | | | | | | | | 51953 |
| | | | | | | | | 51976 |
| | | | | | | | | 51985 |
| | | | | | | | | 51986 |
| | | | | | | | | 51998 |
| | | | | | | | | 52002 |
| | | | | | | | | 52038 |
| | | | | | | | | 57384 |
| | | | | | | | | 60221 |
| | | | | | | | | 60223 |
| | | | | | | | | 60234 |
| | | | | | | | | 60297 |
| | | | | | | | | 71477 |
| | | | | | | | | 49876 |
| | | | | | | | | 49999 |
| *Balistes polylepis* | UW 014876 | --- | 103797 | --- | 893 | 11929 | 73535 | 30333 |
| | UW 025817 | | 103801 | | | | | |
| | | | 103793 | | | | | |
| *Balistes punctatus* | ---- | --- | 102835 | --- | --- | --- | --- |  |
| | | | 103227 | | | | | |
| *Balistes vetula* | ---- | --- | 103907 (a-b) | --- | --- | 30168 | 15638 | 48110 |
| | | | 103863 | | | 11906 | 47961 | |
| | | | | | | 11877 | 52837 | |
| | | | | | | 11878 | 65379 | |
| | | | | | | 32900 | | |
| | | | | | | 11928 | | |
| *Balistoides conspicillum* | ---- | 9484 | 103896 | 38229-12 | 15135 | 15135 | 112946 | 78129 |
| | | | | 54166-1 | | | | |
| *Balistoides viridescens* | ---- | --- | 130942 | --- | --- | 11914 | 110922 |  |
| | | | 113798 | | | 36907 | | |
| | | | 103894 | | | | | |

| | | | | | | | | |
|---|---|---|---|---|---|---|---|---|
| *Canthidermis* | UW 048621 | --- | 108895 | --- | --- | 146629 | 48578 | 17356 |
| *maculata* | | | 15356 | | | 11874 | 61360 | 102516 |
| | | | 103895 | | | 55503 | 74559 | 23252 |
| | | | | | | | 113476 | 20281 |
| | | | | | | | | 49164 |
| | | | | | | | | 49299 |
| | | | | | | | | 49337 |
| | | | | | | | | 49370 |
| | | | | | | | | 49394 |
| | | | | | | | | 49853 |
| | | | | | | | | 49891 |
| | | | | | | | | 49964 |
| | | | | | | | | 49990 |
| | | | | | | | | 57658 |
| | | | | | | | | 60242 |
| | | | | | | | | 60251 |
| | | | | | | | | 60258 |
| | | | | | | | | 60281 |
| | | | | | | | | 60335 |
| | | | | | | | | 30986 |
| *Canthidermis* | | | | | | | | 5032 |
| *sufflamen* | | | | | | | | 27162 |
| | | | | | | | | 49541 |
| | | | | | | | | 49541 |
| | | | | | | | | 49681 |
| | | | | | | | | 49844 |
| | | | | | | | | 49856 |
| | | | | | | | | 49874 |
| | | | | | | | | 49896 |
| | | | | | | | | 71861 |
| | | | | | | | | 49844 |
| | | | | | | | | 57243 |
| *Melichthys niger* | UW 048380 | --- | 103657 | --- | 8674 | 11918 | 91152 | |
| | UW 000546 | | | | | 11919 | | |
| | UW 026434 (a-b) | | | | | | | |
| *Melichthys vidua* | UW 000542 | --- | 131925 (a-b) | --- | --- | 11912 | 16400 | |
| | UW 013045 | | | | | 163091 | 21572 | |
| | | | | | | | 21744 | |
| | | | | | | | 52226 | |
| *Odonus niger* | UW 010542 (a-b) | --- | 103873 | --- | 9822 | 11913 | 19935 | |
| | | | 102779 | | 8326 | 5896 | 21944 | |
| | UW 013061 | | | | | | 31725 | |

| | | | | | | | | |
|---|---|---|---|---|---|---|---|---|
| | | | | | | | 31726<br>110923 | |
| *Pseudobalistes flavimarginatus* | ---- | 12211<br>28375 | 102420 | 35973<br>56845 | --- | 11900<br>24362 | --- |  |
| *Pseudobalistes fuscus* | ---- | --- | 103841 (a-b) | --- | --- | 3770<br>3759<br>6158<br>11893 (a-c) | --- |  |
| *Pseudobalistes naufragium* | ---- | --- | 103800<br>103905<br>100258 | 3449<br>3450 | --- | 41405<br>50737 | --- |  |
| *Rhinecanthus aculeatus* | UW 008237<br>UW 013063<br>UW 013064<br>UW 014387<br>UW 017710 (a-c)<br>UW 018647<br>UW 018649 | --- | --- | ------ | 7184<br>9613 | --- | 47792<br>63522<br>76682<br>89707<br>LW-15 | 79605<br>WCS-915 |
| *Rhinecanthus assasi* | ---- | --- | --- | --- | 20551 | 50082<br>440 | --- |  |
| *Rhinecanthus rectangulus* | UW 013053<br>UW 011096<br>UW 016610 (a-b)<br>UW 026440<br>UW 026439<br>UW un-cataloged (2X) | --- | --- | --- | --- | --- | 44227<br>44229<br>44231<br>55641 | 30485 |
| *Rhinecanthus verrucosus* | UW 009159 | --- | 130753<br>91393 | --- | --- | 24868<br>30516 | 3975<br>52000 | |
| *Sufflamen bursa* | ---- | --- | --- | 42474<br>1245<br>1244 | --- | 5898 | 16399<br>47703<br>63681<br>73656 | 77393 |
| *Sufflamen chrysopterum* | UW 008645 (a-c) | --- | 63882 | --- | 8327 | 8327 | --- | |

| | | | | | | | | |
|---|---|---|---|---|---|---|---|---|
| | UW 009210<br>UW 9379<br>UW 9106 | | | | | | | |
| *Sufflamen fraenatum* | ---- | --- | 105589 (a-c)<br>815933 | --- | --- | 29740 | 73660 |  |
| *Sufflamen verres* | ---- | --- | 100283<br>100251<br>100252<br>103809 | --- | 1095<br>814 | 29628<br>36524<br>45781<br>29716 | --- |  |
| *Xanthichthys auromarginatus* | ---- | 35394 (a-c)<br>10088 (a-b) | 134540<br>134542 (a-c) | 45004 | --- | --- | --- |  |
| *Xanthichthys mento* | ---- | --- | 68601 | 8999-16<br>31778-57<br>31778-60<br>31778-61 | --- | --- | --- |  |
| *Xanthichthys ringens* | ---- | --- | 100989 (a-c)<br>157639 (a-b) | --- | 6247 | 6247<br>11940 | --- | 52104<br>57424 |

**Supplemental Table 2: Habitat classification and data from the primary literature**

| | Reef/ Non-Reef | Flow | Habitat | Reference |
|---|---|---|---|---|
| **ADULT** | | | | |
| *Abalistes stellaris* | Reef | 1 | 10 | Fisher et al. (1990) |
| *Abalistes stellatus* | Reef | 1 | 10 | Bray (2011) |
| *Balistes undulates* | Reef | 3 | 4,6,7,12 | Myers (1991), Bean et al. (2002), Kuwamura (1991) |
| *Balistes capriscus* | Non-Reef | 3 | 4,6,9 | Aiken (1975), Nunoo et al. (2006) |
| *Balistes polylepis* | Non-Reef | 3 | 10,1,11 | Randall and Mundy (1998), Thomson et al. (1979), Kells et al. (2016) |
| *Balistes punctatus* | Non-Reef | 3 | 10,1,11 | Nunoo et al. (2006) |
| *Balistes vetula* | Reef | 3 | 1,2,6,11,12 | Aiken (1975), Cervigón et al. (1992), Lyczkowski-Shultz and Ingram (2003) |
| *Balistoides conspicillum* | Reef | 2 | 6 | Myers (1991), Kulter and Tonozuka (2001) |
| *Balistoides viridescens* | Reef | 3 | 4,6 | Myers (1991), Kuwamura (1991) |
| *Canthidermis maculata* | Non-Reef | 2 | 5,9 | Aiken (1975), Lyczkowski-Shultz and Ingram (2003) |
| *Canthidermis sufflamen* | Non-Reef | 2 | 5,6,8,9 | Aiken (1975), Myers (1991), Lyczkowski-Shultz and Ingram (2003) |
| *Melichthys indicus* | Reef | 2 | 8 | Kulter and Tonozuka |
| *Melichthys niger* | Reef | 3 | 1,6,7,8,12 | Aiken (1975), Myers (1991), Kells et al. (2016) |
| *Melichthys vidua* | Reef | 2 | 6 | Myers (1991), Bean et al. (2002), Kuwamura (1991) |
| *Odonus niger* | Reef | 2 | 6,7 | Myers (1991), Kuwamura (1991) |
| *Pseudobalistes flavimarginatus* | Reef | 1 | 1,4,12 | Myers (1991) |
| *Pseudoblalistes fuscus* | Reef | 3 | 1,4,6,10,12 | Myers (1991), Kuwamura (1991) |
| *Pseudobalistes naufragium* | Non-Reef | 3 | 1,6,12 | Bussing (1995) |
| *Rhinecanthus aculeatus* | Reef | 1 | 1,4,12 | Myers (1991), Kuwamura (1991), Hoover (2003) |
| *Rhinecanthus assasi* | Reef | 1 | 1,10 | Lieske and Myers (1994), Kulter and Tonozuka (2001), |
| *Rhinecanthus lunula* | Reef | 2 | 6 | Bacchet et al. (2006) |
| *Rhinecanthus rectangulus* | Reef | 3 | 1,6,12 | Myers (1991), Kuwamura (1991), Matsuura (2001) |
| *Rhinecanthus verrucosus* | Reef | 1 | 2,4,12 | Myers (1991), Bean et al. (2002), Kuwamura (1991) |
| *Sufflamen albicaudatum* | Reef | 1 | 1 | Myers (1991) |
| *Sufflamen bursa* | Reef | 1 | 6,8,12 | Myers (1991), Bean et al. (2002) |
| *Sufflamen chrysopterum* | Reef | 3 | 1,4,6 | Myers (1991), Bean et al. (2002), Kuwamura (1991) |
| *Sufflamen fraenatum* | Reef | 1 | 1,11,12 | Myers (1991), Kuwamura (1991) |
| *Sufflamen verres* | Reef | 1 | 1,11 | Thomson et al. (1979) |
| *Xanthichthys auromarginatus* | Non-Reef | 2 | 7,6,8 | Kulter and Tonozuka (2001), Hoover (2003) |
| *Xanthichthys lineopunctatus* | Reef | 2 | 6 | Kulter and Tonozuka (2001) |
| *Xanthichthys mento* | Reef | 3 | 8,11 | Kulter and Tonozuka (2001), Hoover (2003), Kells et al. (2016) |

| | | | | |
|---|---|---|---|---|
| *Xanthichthys ringens* | Reef | 2 | 8 | Aiken (1975), Lieske and Myers (1994), Lyczkowski-Shultz and Ingram (2003) |
| **JUVENILE** | | | | |
| *Abalistes stellatus* | | 1 | 9 | Kulter and Tonozuka (2001) |
| *Balistapus undulates* | | 3 | 6,12 | Bean et al. (2002), Kuwamura (1991) |
| *Balistes capriscus* | | 1 | 9 | Longley and Hildebrand (1941), Clements et al. (1991), Hoffmayer et al. (2005), Ballard and Rakocinski (2005) |
| *Balistes polylepis* | | 1 | 1,9 | Thomson et al. (1979), Kells et al. (2016) |
| *Balistes vetula* | | 1 | 9,12 | Robertson (1998) |
| *Balistoides conspicillum* | | 1 | 8 | Kulter and Tonozuka (2001) |
| *Balistoides viridiscens* | | 1 | 1 | Kuwamura (1991) |
| *Canthidermis maculata* | | 1 | 9 | Clements et al. (1991), Hoffmayer et al. (2005) |
| *Canthidermis sufflamen* | | 1 | 9 | Clements et al. (1991), Hoffmayer et al. (2005) |
| *Melichthys vidua* | | 1 | 6,12 | Bean et al. (2002), Kuwamura (1991) |
| *Odonus niger* | | 1 | 1 | Kuwamura (1991) |
| *Pseudobalistes fuscus* | | 1 | 1,9 | Kuwamura (1991) |
| *Rhinecanthus aculeatus* | | 1 | 4 | Kuwamura (1991) |
| *Rhinecanthus assasi* | | 1 | 9,12 | Lieske and Myers (1994) |
| *Rhinecanthus rectangulus* | | 1 | 12 | Kulter and Tonozuka (2001), |
| *Rhinecanthus verrucosus* | | 1 | 3,12 | Bean et al. (2002), Kuwamura (1991) |
| *Sufflamen bursa* | | 1 | 12 | Bean et al. (2002) |
| *Sufflamen chrysopterum* | | 1 | 2,12 | Bean et al. (2002), Kuwamura (1991) |
| *Sufflamen fraenatum* | | 1 | 2 | Kuwamura (1991) |
| *Xanthichthys ringens* | | 1 | 9 | Randall (1968), Clements et al. (1991) |

Flow regimes range from (1) Low flow or sheltered habitat; (2) high flow or open habitat; and (3) mix of low and high/sheltered and open habitat. Habitat types: (1) debris/rubble; (2) sea grass; (3) mangrove; (4) lagoon; (5) epipelagic/pelagic; (6)outer reef; (7) surge zone/strong currents; (8) drop-off; (9) sargassum/floating debris; (10) sand flat; (11) rocky reef; (12) inner reef/sheltered reef.

Supplemental Table 3: Genbank Accessions

| Taxon | *12S* | *16S* | *Rhodopsin* | *Tmo-4C4* | *RAG1* | *BMP4* | *COI* | *CYTB* | *GLYT* | *MYH6* | *RAG2* |
|---|---|---|---|---|---|---|---|---|---|---|---|
| *Abalistes stellaris* | | | | | | KT600846.1 | KF025667.1 | AP009202.1 | | | KT600890.1 |
| *Abalistes stellatus* | AY700248 | AY679632 | EU108845 | EU108823 | AY700318 | | KF025668.1 | | JX190373.1 | | |
| *Acanthaluteres spilomelanurus* | | | | | | | KF025693.1 | KF025759.1 | | | |
| *Acanthaluteres vittiger* | | | | | | | KF025694.1 | KF025760.1 | | | |
| *Acreichthys tomentosus* | | | | | | | KF025695.1 | KF025761.1 | KF139659.1 | | |
| *Aluterus schoepfii* | | | | | | KT600848.1 | KF025698.1 | KF025764.1 | | | |
| *Aluteres heudelotii* | | | | | | | KF025696.1 | KF025762.1 | | | |
| *Aluteres monceros* | | | | | | | KF025697.1 | KF025763.1 | | | |
| *Aluterus scriptus* | | | | | | KT600849.1 | KF025699.1 | KF025765.1 | KF027670.1 | | |
| *Amanses scopas* | | | | | | | KF025700.1 | KF025766.1 | KF027671.1 | KT600891.1 | KT600891.1 |
| *Brachaluteres jacksonianus* | | | | | | | KF025701.1 | KF025767.1 B | | | |
| *Balistapus undulatus* | EU108802 | EU108813 | EU108849 | EU108826 | EU108869 | KT600852.1 | JQ861003.1 | AP009203.1 | KF027662.1 | | KT600894.1 |
| *Balistes capriscus* | AY700238 | AY679622 | EU108846 | EU108824 | AY700308 | KT600853.1 | KF025670.1 | DQ197928.1 | KF139685.1 | | DQ874786.1 |
| *Balistes polylepis* | AY700239 | AY679623 | --- | --- | AY700309 | KT600854.1 | KF025673.1 | KF025738.1 | | | KT600896.1 |
| *Balistes punctatus* | EU108801 | EU108812 | EU108848 | EU108827 | EU108868 | KT600855.1 | KF025674.1 | KF025739.1 | | | KT600897.1 |
| *Balistes vetula* | AY700240 | AY679624 | EU108850 | EU108828 | AY700310 | KT600856.1 | KF025671.1 | AP009204.1 | KF027663.1 | | KT600898.1 |
| *Balistoides conspicillum* | AY700241 | AY679625 | EU108847 | EU108825 | AY700311 | KT600857.1 | KF025672.1 | AP009205.1 | | | KT600899.1 |

| | | | | | | | | | | | |
|---|---|---|---|---|---|---|---|---|---|---|---|
| *Balistoides viridescens* | AY700250 | AY679634 | --- | --- | AY700320 | KT600858.1 | KF025675.1 | KF025740.1 | | | KT600900.1 |
| *Cantherhines fronticinctus* | | | | | | | | | | KT600901.1 | KT600901.1 |
| *Cantherhines pardalis* | | | | | | KT600861.1 | KF025702.1 | KF025769.1 | KF027672.1 | KT600902.1 | KT600902.1 |
| *Cantherhines dumerilii* | | | | | | KT600859.1 | | KF025768.1 | | | |
| *Canthidermis maculata* | AY700242 | AY679626 | EU108851 | EU108829 | AY700312 | KT600862.1 | KF025676.1 | AP009206.1 | | | KT600904.1 |
| *Canthidermis sufflamen* | | | | | | KT600863.1 | | | | | KT600905.1 |
| *Cantherhines pullus* | | | | | | | KF025703.1 | KF025770.1 | JX190377.1 | KT600903.1 | KT600903.1 |
| *Cantherhines sandwichiensis* | | | | | | | | KF025771.1 | | | |
| *Chaetodermis penecilligerus* | | | | | | | KF025704.1 | KF025772.1 | | | |
| *Diodon hystrix* | | | | | | | KF025664.1 | KF025730.1 | | | |
| *Eubalichthys Bucephalus* | | | | | | | KF025705.1 | KF025773.1 | | | |
| *Eubalichthys mosaicus* | | | | | | | KF025706.1 | KF025774.1 | | | |
| *Melichthys indicus* | | | | | | | KF025677.1 | KF025742.1 | | | |
| *Melichthys niger* | AY700243 | AY679627 | EU108852 | --- | AY700313 | KT600865.1 | KF025678.1 | KF025743.1 | | | KT600907.1 |
| *Melichthys vidua* | EU108803 | EU108814 | EU108853 | EU108830 | EU108870 | KT600866.1 | KF025679.1 | AP009207.1 | | | KT600908.1 |
| *Meuschenia hippocrepis* | | | | | | | KF025708.1 | KF025776.1 | | | |
| *Meuschenia trachylepis* | | | | | | | KF025709.1 | KF025777.1 | | | |
| *Meuschenia freycineti* | | | | | | | KF025707.1 | KF025775.1 | | | |
| *Mola mola* | | | | | | | KF025665.1 | KF025731.1 | | | |
| *Monacanthus chinensis* | | | | | | | KF025710.1 | KF025778.1 | KF027673.1 | | |

| | | | | | | | | | | | |
|---|---|---|---|---|---|---|---|---|---|---|---|
| *Monacanthus ciliates* | | | | | | | KF025711.1 | KF025779.1 | | | |
| *Monacanthus tuckeri* | | | | | | | | KF025780.1 | | | |
| *Nelusetta ayraudi* | | | | | | | KF025712.1 | KF025781.1 | | | |
| *Odonus niger* | EU10880<br>4 | EU1088<br>15 | EU108854 | EU108831 | EU108871 | KT600867.1 | KF025680.1 | AP009208.1 | | | KT600909.<br>1 |
| *Oxymonacanthus longirostris* | | | | | | KT600868.1 | KF025713.1 | KF025782.1 | KF027674.1 | KT600910.1 | KT600910.<br>1 |
| *Pergavor janthinosoma* | | | | | | KT600870.1 | KF025719.1 | KF025787.1 | KF139831.1 | KT600912.1 | KT600912.<br>1 |
| *Pergavor melanocephalus* | | | | | | KT600881.1 | KF025720.1 | KF025788.1 | | KT600913.1 | KT600913.<br>1 |
| *Pergavor nigrolineatus* | | | | | | | KF025721.1 | KF025789.1 | KF139832.1 | KT600914.1 | KT600914.<br>1 |
| *Paraluteres prionurus* | | | | | | KT600869.1 | KF025714.1 | KF025783.1 | KF027675.1 | KT600911.1 | KT600911.<br>1 |
| *Paramonacanthus choirocephalus* | | | | | | | KF025715.1 | | | | |
| *Paramonacanthus oblongus* | | | | | | | KF025717.1 | KF025785.1 | | | |
| *Paramonacanthus pusillus* | | | | | | | KF025718.1 | KF025786.1 | | | |
| *Paramonacanthus filicauda* | | | | | | | KF025716.1 | KF025784.1 | | | |
| *Peudomonacanthus peroni* | | | | | | | KF025724.1 | KF025792.1 | | | |
| *Peudomonacanthus macrurus* | | | | | | | KF025723.1 | KF025791.1 | | | |
| *Pseudobalistes naufragium* | | | | | | | | KF025746.1 | | | KT600917.<br>1 |
| *Pseudobalistes flavimarginatus* | EU10880<br>5 | EU1088<br>16 | EU108855 | EU108832 | EU108872 | KT600872.1 | | AP009209.1 | KF139856.1 | | KT600915.<br>1 |
| *Pseudobalistes fuscus* | AY70024<br>4 | AY67962<br>8 | EU108856 | EU108833 | AY700314 | KT600873.1h | KF025681.1 | KF025745.1 | | | KT600916.<br>1 |
| *Rhinecanthus abyssus* | | | | | | KT600875.1 | | | | | KT600918.<br>1 |
| *Pseudobalistes naufragium* | GUO1446<br>4 | GUO144<br>61 | GUO14454 | GUO14453<br>Pending | GUO14460 | KT600874.1 | KF025682.1 | | | | |

| | | | | | | | | | | | |
|---|---|---|---|---|---|---|---|---|---|---|---|
| *Pseudodalutarius nascornis* | | | | | | | KF025722.1 | KF025790.1 | | | |
| *Ranzania laevis* | | | | | | | KF025666.1 | | | | |
| *Rhinecanthus aculeatus* | AY70024 7 | AY67963 1 | EU108857 | EU108834 | AY308790 | KT600876.1 | KF025683.1 | AP009210.1 | KF027664.1 | | KT600919. 1 |
| *Rhinecanthus assasi* | AY70024 5 | AY67962 9 | EU108858 | EU108835 | AY700315 | KT600877.1 | KF025684.1 | KF025748.1 | | | KT600920. 1 |
| *Rhinecanthus lunula* | EU10880 6 | EU1088 17 | EU108859 | EU108836 | EU108873 | | KF025685.1 | KF025749.1 | | | |
| *Rhinecanthus rectangulus* | EU10880 7 | EU1088 18 | EU108860 | EU108837 | EU108874 | KT600878.1 | KF025686.1 | KF025750.1 | | | KT600921. 1 |
| *Rhinecanthus verrucosus* | EU10880 8 | EU1088 19 | EU108861 | EU108838 | EU108875 | KT600879.1 | KF025687.1 | KF025751.1 | KF139869.1 | | |
| *Ranzania laevis* | | | | | | | | KF025732.1 | | | |
| *Rudarius ercodes* | | | | | | | KF025725.1 | | | | |
| *Stephanolepis auratus* | | | | | | | KF025727.1 | | | | |
| *Scobinichthys granulatus* | | | | | | | KF025726.1 | KF025793.1 | | | |
| *Stephanolepis hispidus* | | | | | | KT600880.1 | KF025729.1 | KF025795.1 | KF139893.1 | KT600922.1 | KT600922. 1 |
| *Stephanolepis setifer* | | | | | | KT600881.1 | | KF025796.1 | | KT600923.1 | KT600923. 1 |
| *Stephanolepis cirrifer* | | | | | | | KF025728.1 | KF025794.1 | KF027676.1 | | |
| *Sufflamen albicaudatum* | EU10880 9 | EU1088 20 | EU108862 | EU108839 | EU108876 | | | KF025798.1 | | | |
| *Sufflamen bursa* | AY70024 9 | AY67963 3 | EU108863 | EU108840 | AY700319 | KT600882.1 | | KF025752.1 | | | KT600924. 1 |
| *Sufflamen chrysopterum* | AY70025 1 | AY67963 4 | EU108864 | EU108841 | --- | KT600883.1 | | KF025753.1 | KF139894.1 | | KT600925. 1 |
| *Sufflamen fraenatum* | NC | NC | --- | --- | AY700321 | KT600884.1 | | AP004456.1 | | | KT600926. 1 |

| | | | | | | | | | | | |
|---|---|---|---|---|---|---|---|---|---|---|---|
| | 004416 | 004416 | | | | | | | | | |
| *Sufflamen verres* | GUO14463 | GUO14462 | GUO14455 | GUO14452 | GUO14459 | | KF025688.1 | KF025755.1 | | | |
| *Thamnaconus modestus* | | | | | | | | KF025797.1 | | | |
| *Thamnaconus fajardoi* | | | | | | | | KF025799.1 | | | |
| *Xenobalistes tumidipectoris* | | | | | | | | AP009182.1 | | | |
| *Xanthichthys lineopunctatus* | | | | | | | KF025690.1 | KF025756.1 | | | |
| *Xanthichthys auromarginatus* | AY700246 | AY679630 | EU108865 | EU108842 | AY700316 | KT600887.1 | KF025689.1 | AP009211.1 | | | KT600927.1 |
| *Xanthichthys mento* | EU108810 | EU108821 | EU108866 | EU108843 | EU108877 | KT600888.1 | KF025691.1 | KF025757.1 | | | |
| *Xanthichthys ringens* | EU108811 | EU108822 | EU108867 | EU108844 | EU108878 | KT600889.1 | KF025692.1 | KF025758.1 | KF139910.1 | | KT600928.1 |

**Supplemental Table 4: Best-fit nucleotide substitution models and partitioning strategies identified by PartitionFinder.**

| Subset | Best Model | # of Sites | Contained Partitions |
|---|---|---|---|
| 1 | GTR+I+G | 868 | 12Snonpairing, 16Snonpairing |
| 2 | GTR+I+G | 1120 | 16Spairing, COIp1, 12Spairing, cytbp3 |
| 3 | HKY+G | 1144 | MYH6p1, RAG2p1, RAGp1 |
| 4 | HKY+G | 1979 | BMP4p2, 4c4p1, Glytp1, Rhodp3, RAGp2, MYH6p2, RAG2p2 |
| 5 | K80+G | 490 | RAGp3 |
| 6 | HKY+G | 537 | Rhodp1, Rhodp2, BMP4p1 |
| 7 | HKY+G | 488 | 4c4p2, Glytp2 |
| 8 | K80+G | 487 | Glytp3, 4c4p3 |
| 9 | GTR+I | 160 | BMPp3 |
| 10 | HKY+I+G | 592 | COIp2, cytbp1 |
| 11 | GTR+G | 226 | COIp3 |
| 12 | GTR+G | 365 | cytbp2 |
| 13 | HKY+G | 652 | RAG2p3, MYH6p3 |

**Supplemental Table 5: Taxon coverage of phenotypic data**

| | Life Stage | Fin aspect ratios | Fin incidence angles | Body Shape | Fin Shapes |
|---|---|---|---|---|---|
| *Abalistes stellaris* | Adult | + | + | + | + |
| | Juvenile | + | + | + | + |
| *Abalistes stellatus* | Adult | + | + | + | + |
| | Juvenile | --- | --- | --- | --- |
| *Balistes undulates* | Adult | + | + | + | + |
| | Juvenile | + | + | + | + |
| *Balistes capriscus* | Adult | + | + | + | + |
| | Juvenile | + | + | + | + |
| *Balistes polylepis* | Adult | + | + | + | + |
| | Juvenile | --- | --- | --- | --- |
| *Balistes punctatus* | Adult | + | + | + | + |
| | Juvenile | --- | --- | --- | --- |
| *Balistes vetula* | Adult | + | + | + | + |
| | Juvenile | + | + | + | + |
| *Balistoides conspicillum* | Adult | + | + | + | + |
| | Juvenile | + | + | + | + |
| *Balistoides viridescens* | Adult | + | + | + | + |
| | Juvenile | + | + | + | + |
| *Canthidermis maculata* | Adult | + | + | + | + |
| | Juvenile | + | + | + | + |
| *Canthidermis sufflamen* | Adult | + | + | + | + |
| | Juvenile | + | + | + | + |
| *Melichthys indicus* | Adult | + | + | + | + |
| | Juvenile | --- | --- | --- | --- |
| *Melichthys niger* | Adult | + | + | + | + |
| | Juvenile | + | + | + | + |
| *Melichthys vidua* | Adult | + | + | + | + |
| | Juvenile | --- | --- | --- | --- |
| *Odonus niger* | Adult | + | + | + | + |
| | Juvenile | + | + | + | + |
| *Pseudobalistes flavimarginatus* | Adult | + | + | + | + |
| | Juvenile | + | + | + | + |
| *Pseudoblalistes fuscus* | Adult | + | + | + | + |
| | Juvenile | + | + | + | + |
| *Pseudobalistes naufragium* | Adult | + | + | + | + |
| | Juvenile | + | + | + | + |
| *Rhinecanthus aculeatus* | Adult | + | + | + | + |

| | | | | | |
|---|---|---|---|---|---|
| | Juvenile | + | + | + | + |
| *Rhinecanthus assasi* | Adult | + | + | + | + |
| | Juvenile | + | + | + | + |
| *Rhinecanthus lunula* | Adult | + | + | + | + |
| | Juvenile | --- | --- | --- | --- |
| *Rhinecanthus rectangulus* | Adult | + | + | + | + |
| | Juvenile | + | + | + | + |
| *Rhinecanthus verrucosus* | Adult | + | + | + | + |
| | Juvenile | + | + | + | + |
| *Sufflamen albicaudatum* | Adult | + | + | + | + |
| | Juvenile | + | + | + | + |
| *Sufflamen bursa* | Adult | + | + | + | + |
| | Juvenile | + | + | + | + |
| *Sufflamen chrysopterum* | Adult | + | + | + | + |
| | Juvenile | + | + | + | + |
| *Sufflamen fraenatum* | Adult | + | + | + | + |
| | Juvenile | + | + | + | + |
| *Sufflamen verres* | Adult | + | + | + | + |
| | Juvenile | --- | --- | --- | --- |
| *Xanthichthys auromarginatus* | Adult | + | + | + | + |
| | Juvenile | --- | --- | --- | --- |
| *Xanthichthys lineopunctatus* | Adult | + | + | + | + |
| | Juvenile | + | + | + | + |
| *Xanthichthys mento* | Adult | + | + | + | + |
| | Juvenile | --- | --- | --- | --- |
| *Xanthichthys ringens* | Adult | + | + | + | + |
| | Juvenile | + | + | + | + |

Presence/absence of phenotypic trait data between adult and juvenile triggerfishes. Fin aspect ratios = both dorsal and anal fin aspect ratios, fin incidence angles = the presence of incidence angles for both the anal and dorsal fins, fin shapes = shape data for both the anal and dorsal fins. + = presence of data; --- = absence of data.

**Supplemental Table 6: Percent variation captured by each relative warp axis**

| | *Anal Fin* | *Dorsal Fin* | *Body Shape* |
|---|---|---|---|
| RelWarp1 | 38.25 | 49.8 | 41.2 |
| RelWarp2 | 35.24 | 24.3 | 20 |
| RelWarp3 | 15.74 | 13.8 | 13.2 |

**Figure S1**: Landmarks and measurements used for analyses. (A) Landmarks (dark) and sliding semi-landmarks (light) used for shape analyses of the body. (B) Graphical representation of landmarks (dark) and sliding semi-landmarks (light) used for shape analyses of the fin and areas measured to quantify the emergent mechanical properties of the fins. Numbers correspond to points discussed in the primary text.

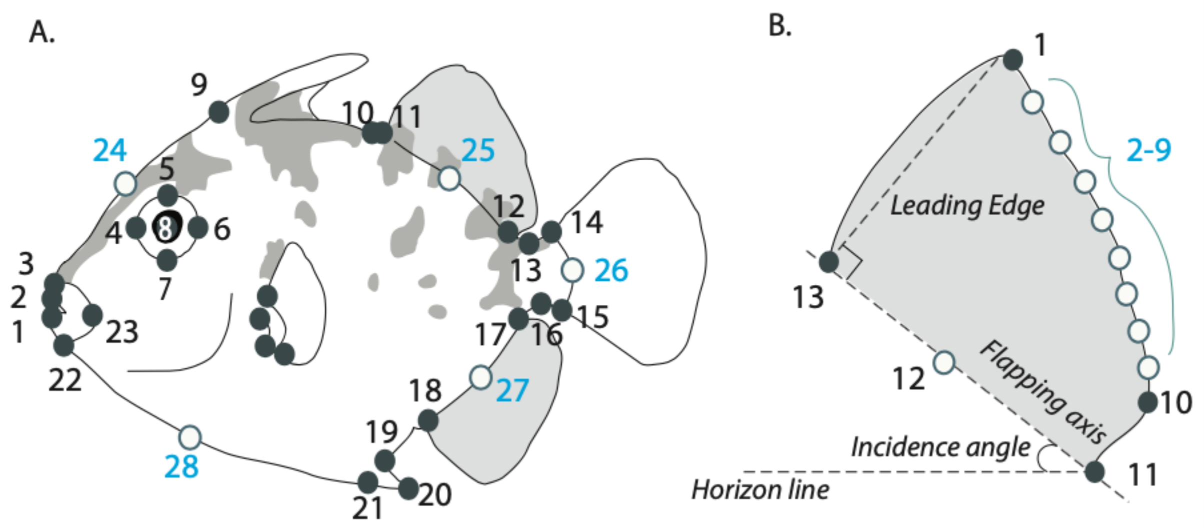

**Figure S2:** Divergence time estimates of Balistidae. (A) Comparison of divergence time estimated using all loci (left) versus estimates based on loci filtered by (B) PI profile shape. (C) Comparison of branch length 95% HPD intervals between the two datasets. (D) Comparison of age estimates between four data inclusion and partitioning strategies and the prior expectation for crown triggerfishes, filefishes, and the most recent common ancestor of triggerfishes and pufferfishes.

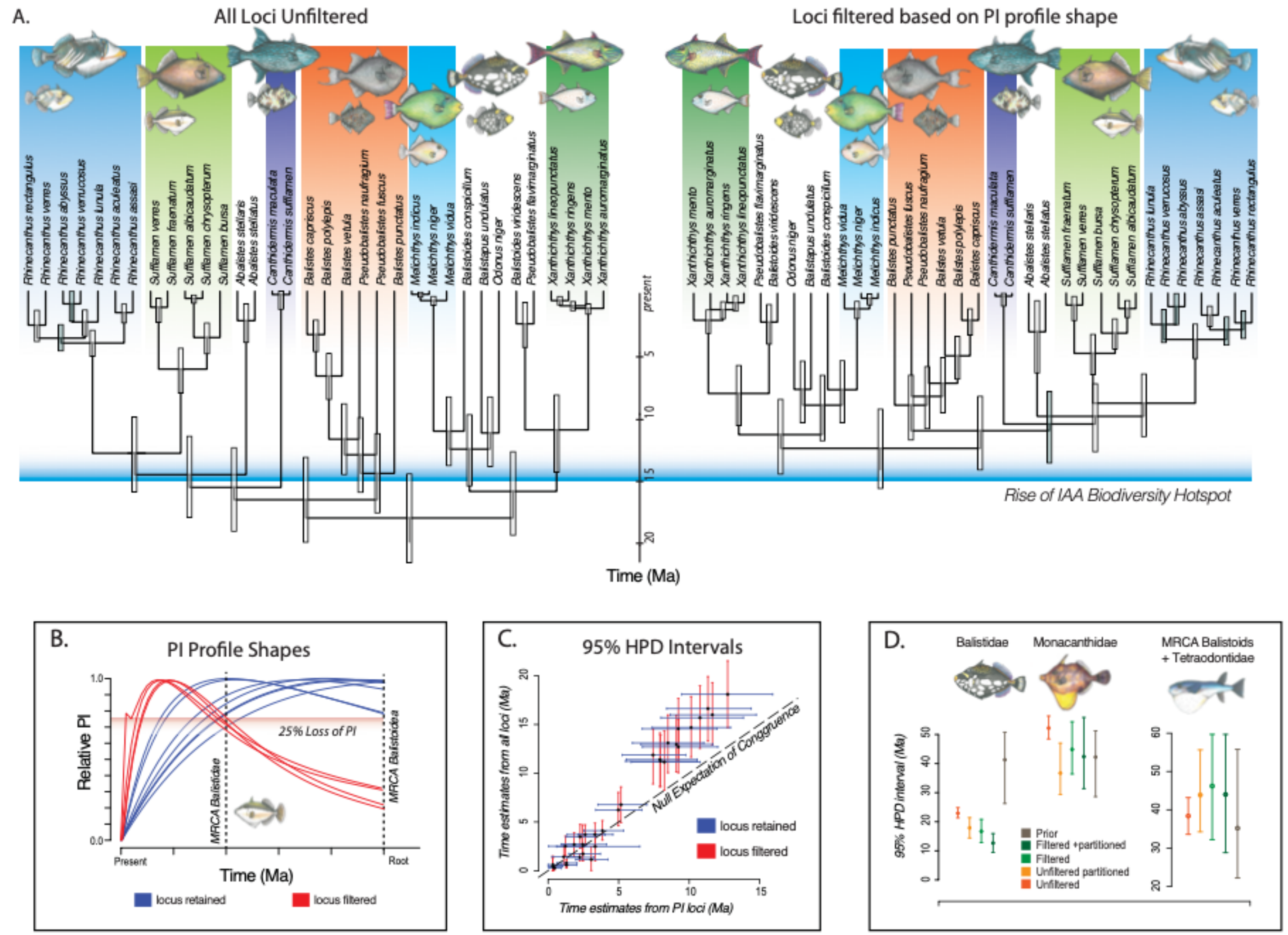

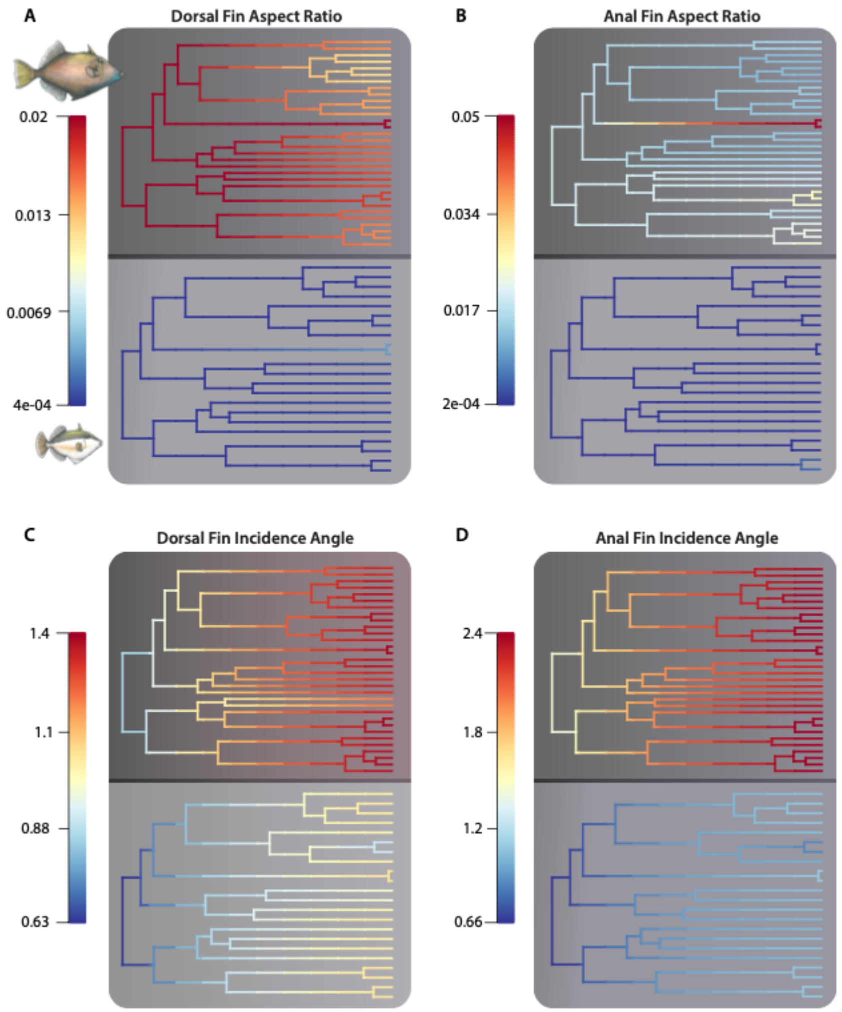

**Figure S3.** Visualization of rates of trait diversification between adults (top) and juveniles (bottom) for A) dorsal fin aspect ratio; B) anal fin aspect ratio; C) dorsal fin incidence angle; and D) anal fin incidence angle.
